# Oxidative stress promotes axonal atrophy through alterations in microtubules and EB1 function

**DOI:** 10.1101/2024.07.12.603221

**Authors:** Samuel Shields, Oliver Wilkes, Illana Gozes, Natalia Sanchez-Soriano

## Abstract

Axons are crucial for transmitting neurochemical signals. As organisms age, the ability of neurons to maintain their axons declines; hence aged axons are more susceptible to damage or dysfunction. Understanding what causes axonal vulnerability is crucial for developing strategies to enhance overall resilience of neurons, and to prevent their deterioration during ageing or in age-related neurodegenerative diseases.

Increasing levels of reactive oxygen species (ROS) causes oxidative stress, a hallmark of ageing and age-related diseases. Despite this association, a causal relationship between oxidative stress and neuronal ageing remains unclear, particularly how subcellular physiology is affected by ROS.

By using *Drosophila*-derived primary neuronal cultures and a recently developed *in vivo* neuronal model of ageing, which involves the visualisation of *Drosophila* medulla neurons, we investigated the interplay between oxidative stress, neuronal ageing and the microtubule cytoskeleton. We find that oxidative stress as a key driver of axonal and synaptic decay, including the appearance of axonal swellings, microtubule alterations in both axons and synapses and the morphological transformation of axonal terminals during ageing. We demonstrate that increased ROS sensitises the microtubule plus end binding factor, end-binding protein 1 (EB1), leading to microtubule defects, affecting neuronal integrity. Furthermore, manipulating EB1 proved to be a valuable therapeutic strategy to prevent ageing hallmarks observed in conditions of elevated ROS. In summary, we demonstrate a mechanistic pathway linking cellular oxidative stress, the microtubule cytoskeleton and axonal deterioration during ageing and provide evidence of the therapeutic potential of enhancing microtubule plus end physiology to improve the resilience of axons.

## Introduction

Ageing is associated with a decline in cognition, motor and sensor function, impacting activities of daily living and quality of life (Strawbridge et al., 2000). The determinants and cellular processes of ageing in the brain are not well understood, highlighting the need for in depth study of this process.

Reactive oxygen species (ROS) are molecules produced as a by-product of respiration with the capacity to be deleterious in high concentrations. An array of molecular (e.g., glutathione [GSH], vitamin E) and enzymatic (e.g., superoxide dismutases [SODs], catalase) antioxidants act to control ROS levels throughout cells. However, increasing levels of ROS (oxidative stress) classically correlate with age in humans and animal models (Choksi et al., 2007, Albrecht et al., 2011, Bouzid et al., 2014, Lennicke and Cochemé, 2020, Rizvi et al., 2021). Oxidative stress is also implicated in the pathology of various neurodegenerative diseases, including Alzheimer’s disease (AD), Parkinson’s disease (PD) and amyotrophic lateral sclerosis (ALS) (Liu et al., 2017). Recent studies demonstrate that ROS are akin to second messengers, capable of regulating many downstream signalling functions. During ageing, oxidative stress may perturb redox signalling and homeostasis and therefore negatively impact cellular function (Jones, 2015, Lennicke and Cochemé, 2020). While this redox theory of brain ageing has gained support in recent years, the specific cellular and molecular mechanisms through which oxidative stress affects neurons are not understood.

Microtubules (MTs) are filamentous polymers, comprising non-covalent longitudinal arrangements of α- and β-tubulin heterodimers, and are a key constituent of the cytoskeleton (Hahn et al., 2019). MT integrity is required for the proper functioning of the nervous system. In axons, MTs are organised in densely packed parallel bundles, and act as key proponents of axonal structural resistance and morphology by influencing axon calibre (Friede and Samorajski, 1970, Prokop, 2020, Okenve-Ramos et al., 2024). MTs also facilitate the transport of essential material, including organelles, RNAs, proteins and synaptic vesicles in neurons (Wittmann and Waterman-Storer, 2001, Bodaleo and Gonzalez-Billault, 2016). Dysregulation of the MT cytoskeleton has been reported in the ageing brain in human pyramidal neurons and in axons of aged Resus monkey brains (Cash et al., 2003, Fiala et al., 2007). Our group has shown that similar alterations in MT density and organisation are found in medulla neurons situated in the *Drosophila* brain, and that these changes are a true hallmarks of ageing in neurons (Okenve-Ramos et al., 2024). Given that perturbed MT function is also linked to a plethora of age-related neurodegenerative diseases, MT deterioration could represent a key event driving neuronal vulnerability in brain ageing and neurodegenerative conditions. Yet, what causes such alterations on MTs in neurons and their axons during ageing, lacks comprehensive exploration.

*In vitro* studies of myocytes revealed that increased ROS can induce alterations in the properties of MTs (Landino et al., 2007, Drum et al.,2016). This observation led us to hypothesise that neuronal deterioration in ageing could be propelled by oxidative stress, adversely affecting the function of MTs. To test this, we used a previously described model to study the cell biology of neuronal ageing in the *Drosophila* optic lobe. We found that chemical inducers of oxidative stress or genetic reduction of enzymatic (SOD1 and SOD2) and non-enzymatic antioxidants (GSH) exacerbates ageing hallmarks, including the appearance of axonal swellings and bulbous synaptic terminals. We also showed that enhancement of the antioxidant response in neurons by manipulating kelch-like ECH-associated protein 1 (Keap1) levels, a redox-sensitive direct inhibitor of nuclear factor erythroid 2-related factor 2 (NRF2), attenuates ageing hallmarks. Importantly, we discovered that oxidative stress induces MT dysfunction *in vitro* and *vivo*, suggesting a key role during the process of ageing. Furthermore, we showed that changes in MT organisation, mediated through high levels of ROS, are caused by impaired end-binding protein 1 (EB1) activity, and that enhancing EB1 is successful in preventing ROS-induced, age-related neuronal deterioration. This new understanding of the intricate interplay between ageing, oxidative stress and the MT cytoskeleton may offer novel therapeutic strategies to safeguard neuronal integrity as individuals age.

## Materials and Methods

### Fly stocks and husbandry

Flies were maintained on standard fly food containing cornflour, glucose, yeast and agar. For experiments involving ageing, flies were maintained at 25°C during development and transferred to 29°C post-eclosion to expedite the ageing process. Flies were maintained at low densities (maximum of 20 flies per vial) and were transferred to new vials with fresh food every 3 days. Both males and females were used for ageing experiments. Flies within each vial were all of the same sex and all analytical comparisons were age and sex matched accordingly.

The Gal4 driver lines used in this study were *elav-Gal4* (3^rd^ chromosomal, expressing pan-neuronally at all stages (Luo et al., 1994)) and *GMR31F10-Gal4* (Bloomington stock 49685; expressing in T1 medulla neurons; (Qu et al., 2019)). Mutant alleles used in this study were: *SOD1^n1^* (Phillips et al., 1995); (Bloomington #24492), *SOD1^n64^* ((Phillips et al., 1995); Bloomington #7451), *SOD2^Δ02^* (Bloomington #27643) and *EB1^04524^* (Elliott et al., 2005). Lines used for overexpression experiments were *UAS-GFP-α-tubulin84B* (Grieder et al., 2000), *UAS-myr::tdTomato* (Bloomington #32222), *UAS-catalase* (Missirlis et al., 2001), *UAS-EB1::GFP* and *UAS-EB1::mCherry* (Alves-Silva et al., 2012).

### Adult drug feeding

PQ (1,1′-dimethyl-4,4′-bipyridinium dichloride; Merck) and DEM (diethyl (*Z*)-but-2-enedioate; Alfa Aesar) feeding protocols were adapted from (Niveditha et al., 2017). Flies were maintained for approximately 14 days (middle aged) prior to drug feeding. Flies were then transferred to empty food vials containing layered 188 mm Whatman™ filter paper soaked with 500 mL drug/vehicle diluted in 2.5% sucrose (Fisher)/Dulbecco’s phosphate-buffered saline (Sigma, RNBF2227) solution. Flies were exposed to the drug for 24 hours before being transferred back to standard fly food for a further 24 hours (without the drug). This 24-hour cycle is then repeated until flies had received a total of four 24-hour doses (over an 8-day period). Flies were then prepared for brain dissections and confocal imaging (*described below*).

### Live adult brain dissections and microscopy

*Drosophila* brains were dissected in Dulbecco’s PBS after initial anaesthetisation on ice for 1.5 minutes. This method of anaesthetisation has been shown to not to affect MT physiology in T1 cells (Okenve-Ramos et al., 2024). Dissected brains were mounted in Dulbecco’s PBS on MatTek glass-bottom dishes (P35G1.5-14C), with a spacer and coverslip. Brains were then immediately imaged using a 3i Marianas Spinning Disk with a 63× 1.4 NA Zeiss Plan Apochromat lens and FLASH4 sCMOS (Hamamatsu) camera (incubated at 26°C); z-stacks of whole brains were taken using a slice interval of 0.3 µM. All imaging of adult brains was conducted at the Centre for Cell Imaging at the University of Liverpool. For each experimental repeat, sex was kept consistent and data were normalised to respective controls.

### Brain image analyses

Using FIJI/ImageJ, several maximum projection images were generated from captured z-stacks to aid accurate visualisation and analysis of layered T1 axons and axon terminals. For consistency, analyses of axonal membranes (e.g. membrane swellings) and MT-related phenotypes were restricted to the same stretch of axon bundles. The number of axonal swellings and regions of disorganised MTs were recorded and presented per nerve bundle.

For axon terminal analyses, maximum projection images of the terminal compartments were generated from the obtained z-stacks. A 70 µm × 70 µm area is outlined in the centre of each medulla, where most axon terminals are easily identifiable and will be evaluated (this included approximately 30–50 synaptic terminals per medulla). The proportion of axon terminal swellings and terminals with fragmented/broken MTs per medulla and the mean number of synaptic MTs per terminal were quantified. For qualitative measurements (i.e. swollen and fragmented phenotypes), data were presented as a percentage of all terminals evaluated per medulla. For axonal diameter, maximum projections were generated and oriented to view axons in horizontal parallel alignment to facilitate consistency in the analysis. Images containing only the membrane channel were processed using the ‘tubeness’ plugin available within FIJI/ImageJ. To eliminate biases in measuring the diameter of axonal membranes, a 10 µm × 10 µm grid was overlayed and measurements were obtained using the Fiji/ImageJ line tool at points where T1 neurons overlap with the overlaid grid. These values were used to calculate the mean diameter per axon.

### Primary neuronal cell culture and drug treatments

*Drosophila* primary neuronal cultures were carried out using methods previously described (Prokop et al., 2012, Voelzmann and Sanchez-Soriano, 2022).

Drug treatments were carried out at 0–3 DIV, as stated throughout according the experimental approach. Compounds used were: 100 µM PQ (vehicle: H2O), 100 µM–1 mM DEM (vehicle: ethanol), 100 µM Trolox ([±]-6-Hydroxy-2,5,7,8-tetramethylchromane-2-carboxylic acid; Merck), 100 µM nocodazole and 1 nM NAP (a gift from Illana Gozes, Tel Aviv University, Israel).

### Immunocytochemistry of cultured neurons

Unless otherwise stated, cultured neurons were fixed in 4% paraformaldehyde (PFA) in 0.05 M phosphate buffer (pH 7.0–7.2) for 30 minutes at room temperature. For mitochondrial staining, primary neurons were washed with fresh sterile *Schneider’s* medium and incubated with 40 nM MitoTracker Red CMXRos (Invitrogen) for 30 minutes at room temperature, prior to fixation with 4% PFA, followed by immunohistochemistry staining (*see below*). For analyses of EB1 comets, primary *Drosophila* neurons were fixed in ice-cold (stored at –80°C) +TIP-fixative solution (90% methanol, 3% formaldehyde, 5 mM sodium carbonate, pH 9; (Hahn et al., 2021, Rogers et al., 2002)) for 10 minutes. Antibody staining and washes were performed with PBT (PBS supplemented with 0.3% Triton X-100).

Primary antibodies used in this study were: Cy3-751 conjugated anti-HRP (goat, 1:100; Jackson ImmunoResearch), anti-α-tubulin (mouse, 1:1000; T9026, Merck), anti-DmEB1 (rabbit, 1:2000, gift from H. Ohkura). Secondary antibodies used were FITC-, Cy3- or Cy5-conjugated secondary antibodies (goat, 1:100; Jackson ImmunoResearch). Cells were embedded in Vectashield (H-1000, VectorLabs).

### Microscopy of cultured neurons and image analyses

Fixed *Drosophila* cells were captured using a Nikon eclipse 90i with a Retiga 3000 camera (QImaging) or a 3i Marianas Spinning Disk using a 63× 1.4 NA Zeiss Plan Apochromat lens and FLASH4 sCMOS (Hamamatsu) camera at the Centre for Cell Imaging at the University of Liverpool. To measure the degree of MT disorganisation or curling in axons, a ‘MT-disorganisation index’ (MDI) was used. Using FIJI/ImageJ, the areas of disorganised MTs per neuron were measured using the freehand selection tool. The total area of MT disorganisation was then divided by the respective axon length, (measured using the segmented line tool), and multiplied by 0.5 µm (average axonal diameter) to account for the expected axonal area exempt of disorganised or unbundled MTs (Prokop et al., 2012, Voelzmann and Sanchez-Soriano, 2022). EB1 comet lengths were measured using the FIJI/ImageJ line tool and data were presented as mean EB1 comet length per cell.

### Statistical analyses

Statistical analyses were performed in GraphPad Prism 9; statistical tests are stated in the relevant figure legends. Exact p-values are indicated in graphs. Sample sizes and other statistical values are indicated in figure legends.

## Results

### Reductions in SOD activity or GSH levels induces microtubule defects and exacerbates axonal and synaptic ageing hallmarks

To evaluate the impact of ROS on MTs and neuronal ageing *in vivo*, we used a novel *Drosophila* model, previously established by our group (Okenve-Ramos et al., 2024). This model allows the study of the cell biology of ageing in T1 medulla neurons. T1 neurons represent a neural subtype residing in the medulla of the *Drosophila* optic lobe, with cell bodies localised at the outer medulla cortex. The T1 medulla axon, which terminates and synapses within the M2 layer, is easily resolved and has been previously subject to evaluation during ageing (Okenve-Ramos et al., 2024). Alterations in axonal (swellings, reduced diameter) and synaptic morphology (swellings and fragmentation) and MT defects, such as foci with unbundled disorganised MTs and reductions in the number of MTs extending into synaptic terminals, referred to as hallmarks of ageing, develop over time, and are linked to ageing appearing within a few weeks of *Drosophila* life in T1 neurons (Okenve-Ramos et al., 2024).

SODs are metalloenzymes which catalyse the conversion of superoxide radicals to hydrogen peroxide, which is then further reduced to water by peroxidases. SOD1 is predominantly cytosolic, whereas SOD2 is localised to mitochondria. We utilised two amorphic alleles of SODs, *SOD1^n64^* and *SOD2^Δ02^* (Phillips et al., 1995, Zheng and Sehgal, 2009). Since both alleles cause adult lethality in homozygosis, their effects in the T1 ageing model were studied in heterozygosis. In this model, MT cytoskeletons were labelled with *UAS-α-Tubulin84B::GFP* and cell membranes with *UAS-myristoylated-tomato,* under the control of the T1-specific Gal4 driver, *GMR31F10-Gal4* (Okenve-Ramos et al., 2024).

We found that *SOD2^Δ02^*^/+^ flies aged 30–34 days displayed a significant, 3-fold increase in the frequency of unbundled/disorganised MTs within axons as well as a significant increase in the number of axonal swellings (**Figure 1**). *SOD1^n64/+^* failed to significantly enhance axon swellings or MT unbundling despite increasing trends. Since the expression of one wildtype copy of SOD1 may provide sufficient function, we tested T1 neurons-specific knockdown of SOD1, using a validated SOD1 RNAi strain (*^SOD1GL01016^* (Chaplot et al., 2019)); SOD1 knockdown significantly increased the frequency of MT unbundling and axonal swellings in aged specimens (**Figure 2**).

**Figure 1.**
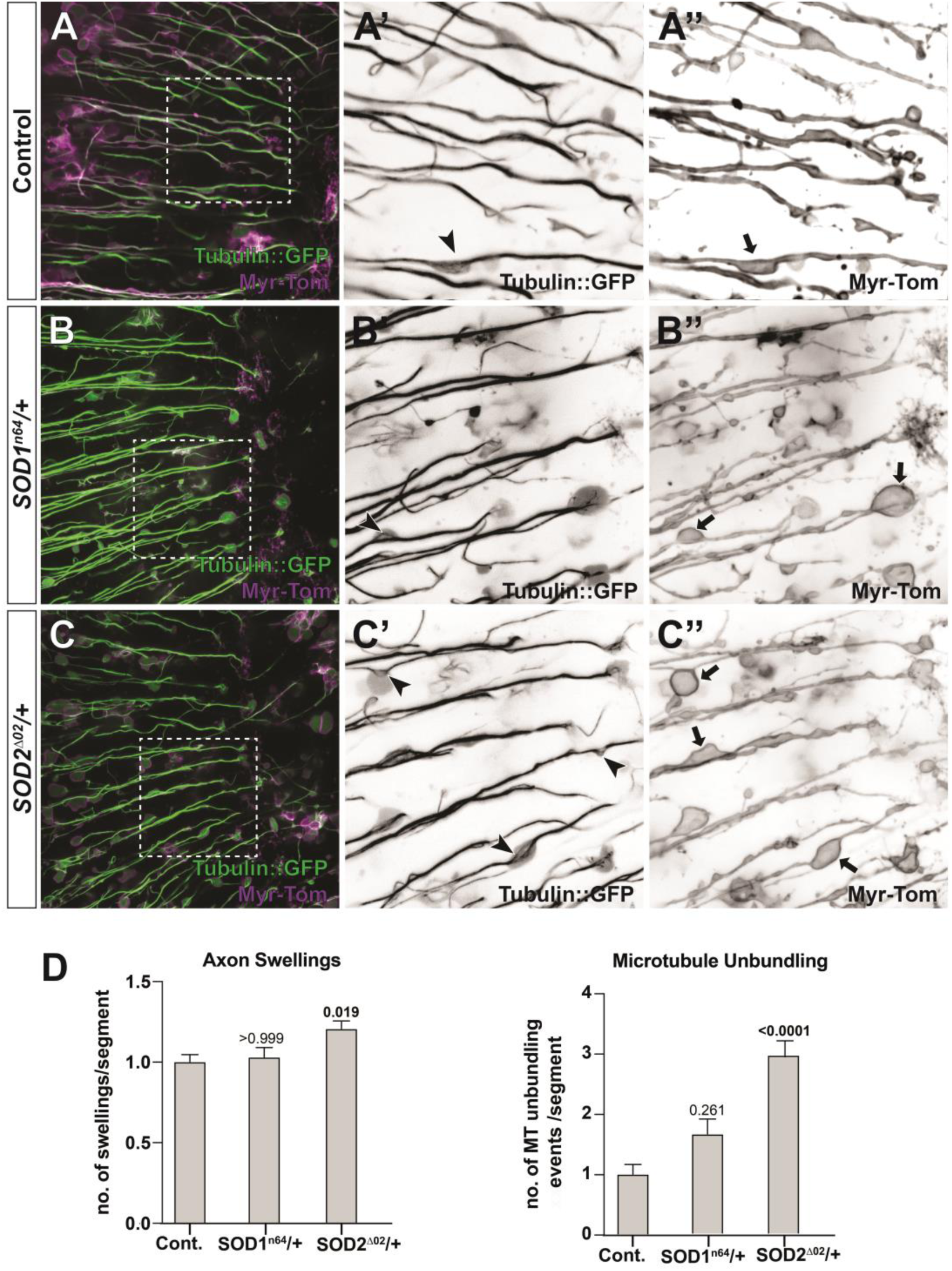
Mutant SOD2 exacerbates neuronal ageing hallmarks of axonal swellings and microtubule disorganisation in brains from aged *Drosophila*. (A–C’’) Medulla regions of adult brains aged 31–35 days post eclosure, which show T1 axons labelled with GFP-tagged α-tubulin (tubulin::GFP) and the plasma membrane marker myristoylated-Tomato (myr-Tom). Magnified inverted-grayscale images of areas outlined by dashed white boxes in A, B, and C are, respectively, shown for tubulin::GFP (A’, B’, C’) and Myr-Tom (A’’, B’’, C’’). Aged neurons in the absence (A) or presence of *SOD1^n64/+^* (B) or *SOD2^Δ02/+^* (C) mutant backgrounds are compared. *SOD2^Δ02/+^* enhance phenotypes in ageing neurons, comprising axon swellings (black arrows) and MT unbundling (arrow heads). (D) Quantitative analyses of the frequency of axonal swellings and microtubule unbundling; bars represent normalised mean ± SEM; p values are shown above each bar, as assessed by Kruskal*-*Wallis one-way tests. Data were collated across four individual repeats, from a minimum of 355 axonal segments from 32 control medullas, 26 *SOD1^n64/+^* medullas and 33 *SOD2^Δ02/+^* medullas. Scale bars = 10 μm.

**Figure 2.**
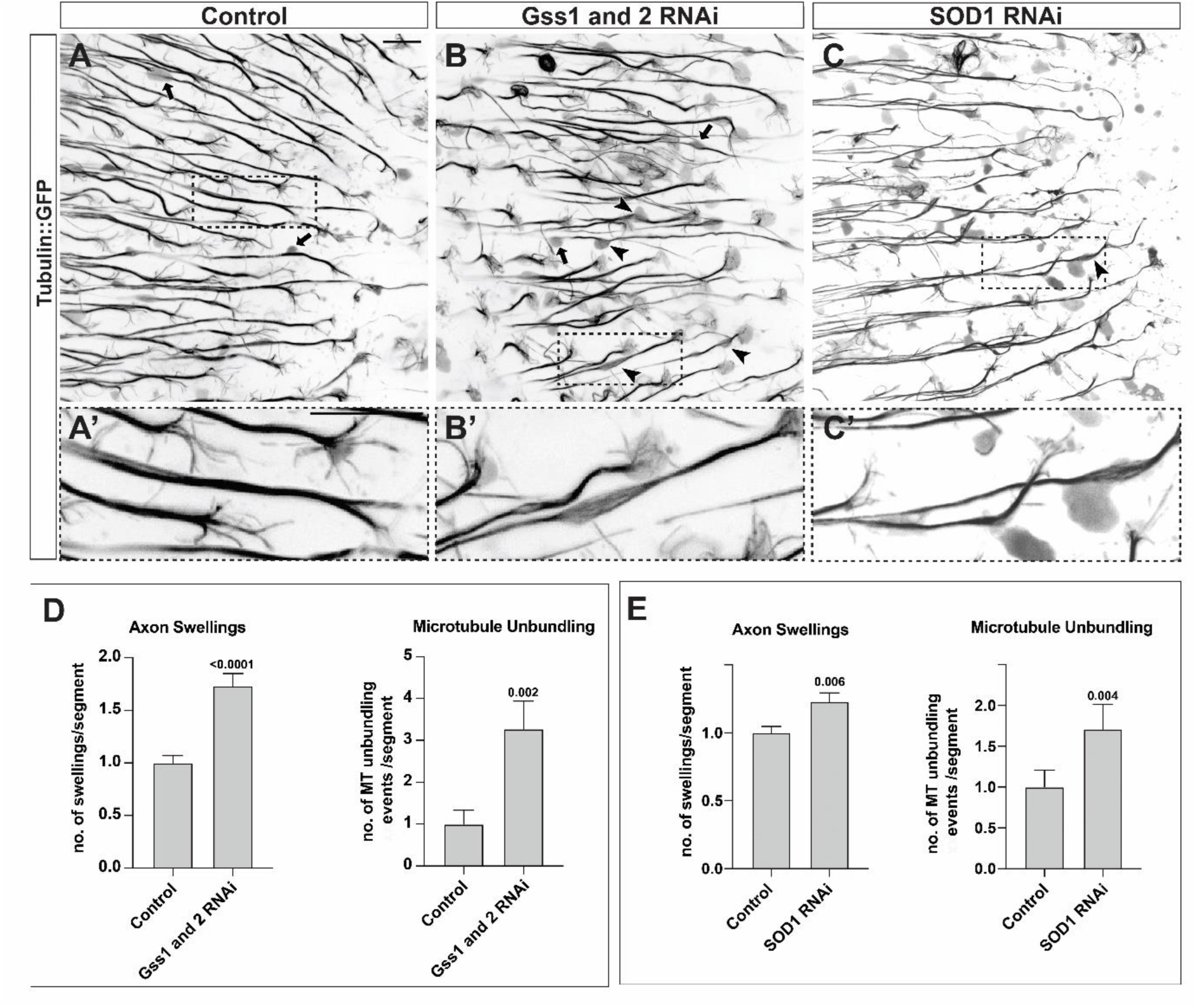
Knocking down *Gss1 and 2* and *SOD1*, enhances age-related axonal swellings and microtubule unbundling. (A–C’) Medulla regions of adult brains aged 30–33 days post eclosure, which show T1 axons labelled with GFP-tagged α-tubulin (tubulin::GFP) presented as inverted greyscale images. Aged neurons in the absence (A) or presence of *Gss1 and 2* knockdown (*Gss1 and 2 RNAi;* B) or *SOD1* knockdown (*SOD1-RNAi*; C) are compared. *Gss1 and 2 and SOD1* knockdown, enhance phenotypes in ageing neurons, comprising axon swellings (arrows) and MT unbundling (arrow heads); dashed boxed areas are magnified and shown in A’, B’, C’. (D and E) Quantitative analyses of the frequency of axonal swellings and microtubule unbundling for Gss1 and 2 (D) and SOD1 (E) knockdown; bars represent normalised mean ± SEM; p values are shown above each bar, as assessed by Mann-Whitney rank sum tests. Data were collated across three individual repeats; from a minimum of 304 axonal segments from 19 control and 19 Gss1 and 2 RNAi medullas in D and 23 control and 18 SOD1 RNAi medullas in E. Scale bars = 10 μm.

Age-related changes in T1 morphology is evident at synaptic terminals, including the development of swollen or bulbous membranes, that can appear fragmented; reductions in the number of invading MTs at synapse have also been reported in T1 neurons with age (Okenve-Ramos et al., 2024). Given that SOD2 had a stronger impact on axonal ageing, we further assessed whether similar effects were observed at the level of synapses. Supporting our axonal findings, *SOD2^Δ02/+^* flies had significantly higher proportions of swollen and fragmented membranes, and significantly fewer MTs visible in synapses (**Figure 3**).

**Figure 3.**
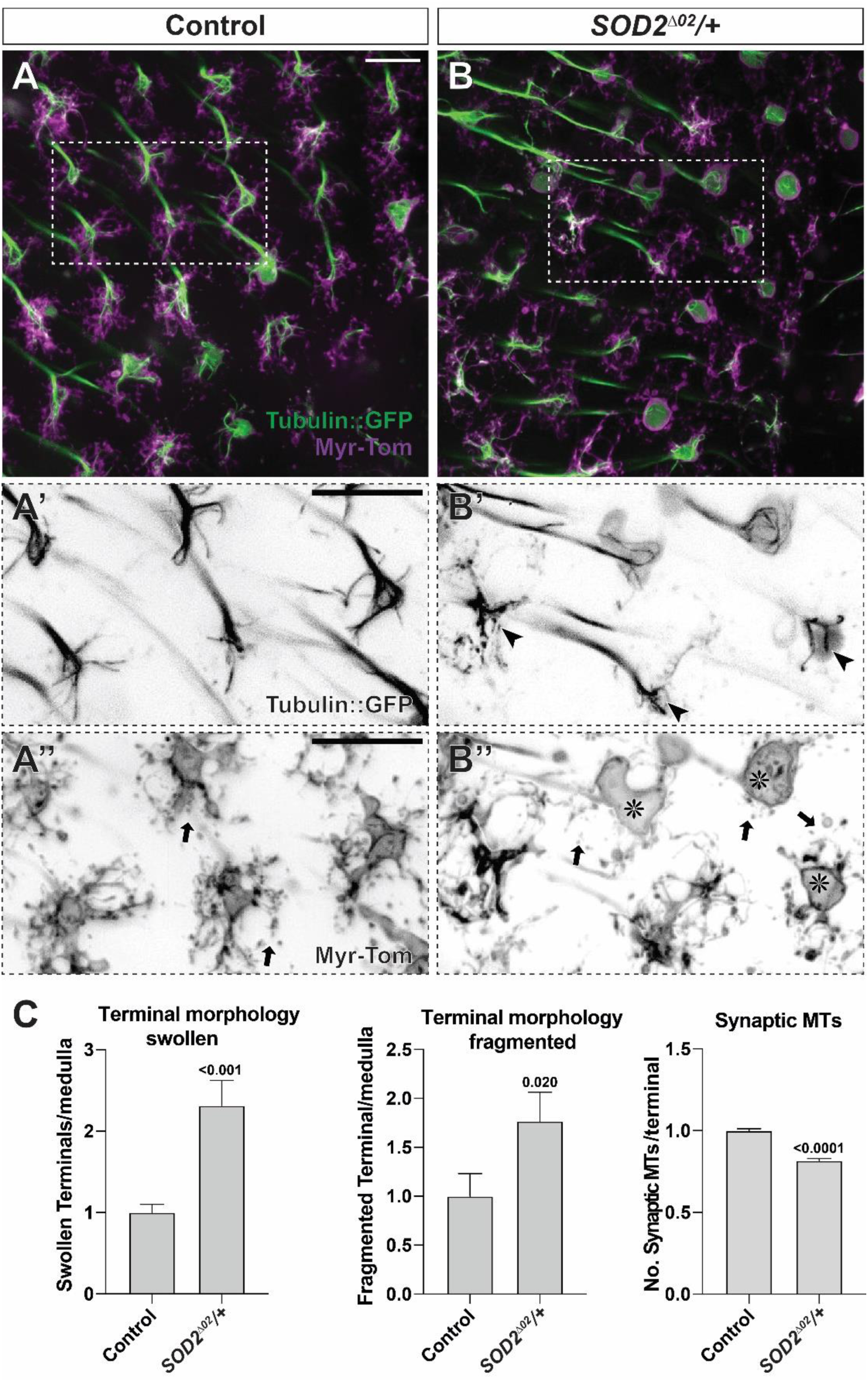
Mutant SOD2 induces a decrease in synaptic microtubules and worsens the deterioration of T1 synaptic terminals in brains of aged *Drosophila*. A–B’’) Axon terminals of T1-cell projections in the medulla of aged *Drosophila* (31–35 days post eclosure), labelled with GFP-tagged α-tubulin (tubulin::GFP) and the plasma membrane marker myristoylated-Tomato (myr-Tom). Magnified inverted-greyscale images of areas outlined by dashed boxes in A, B are, respectively, shown for tubulin::GFP (A’, B’) and Myr-Tom (A’’, B’’). Ageing phenotypes at the synaptic terminals, including an increase in swollen (asterisks) and broken terminals (arrows), and a decrease in synaptic MTs (arrowheads), are enhanced in the presence of *SOD2^Δ02/+^* mutant backgrounds. (C) Quantitative analyses of the frequency of swollen and fragmented terminals and the number of synaptic microtubules in control and *SOD2^Δ02/+-^*mutant backgrounds; bars represent normalised mean ± SEM; p values are shown above each bar, as assessed by Mann-Whitney rank sum tests. Data were collated across four individual repeats; from around 700 synapses from 31 control and 31 *SOD2^Δ02/+^* medullas. Scale bars = 10 μm.

Another layer of the antioxidant defence in cells is provided by small molecule, or non-enzymatic, antioxidants, which function as ROS scavengers or cofactors for ROS detoxification and metabolism. An integral redox couple and antioxidant in cells is GSH. GSH synthetase (Gss) is an essential enzyme in *de novo* synthesis of GSH. We hypothesised that reducing GSH levels, via Gss1 and Gss2 knockdown, may impact physiological ageing within the T1 model. Like the results observed with SOD2/SOD1, knockdown of Gss1 and Gss2 significantly exacerbated the frequency of MT unbundling and axonal swellings, compared with age-matched controls (**Figure 2A, B and D**). Therefore, these data demonstrate that diminishing antioxidant activity can alter basic MT properties in axons, and exacerbate age-related changes in axon and synaptic morphology, by cell autonomous mechanisms.

### Enhancing antioxidant responses by knockdown of Keap1 protects MT deterioration and attenuates axonal and synaptic decay during ageing

Next, we investigated whether enhancing the antioxidant defence system could alleviate negative consequences of ageing on neuronal MTs, and axon and synaptic morphological. To gain this insight, we used two approaches: overexpressing a single enzymatic antioxidant, catalase, which is important for the elimination of H_2_O_2_ in cells (Deisseroth and Dounce, 1970, von Ossowski et al., 1993); or promoting the global network of antioxidant and redox signalling pathways by knocking Keap1, a repressor of NRF2 (Suzuki et al., 2019).

Flies overexpressing catalase had significantly more axonal swellings than aged-matched controls (31– 34 days), and there was no change in the frequency of MT unbundling (**Figure S1**). However, this may suggest that a fine balance between functional ROS levels and oxidative stress may be necessary. To further understand these results, we tested if catalase would have beneficial effects in conditions of exacerbated levels of ROS. To achieve this, we optimised a feeding protocol for the delivery of the bipirydyl redox cycler, paraquat (PQ), which promotes the formation of mitochondrial superoxide radicals (Cochemé and Murphy, 2008, Mitchell et al., 1983). Flies were raised on standard food for 14– 16 days, prior to exposure with PQ or vehicle dissolved in 2.5% sucrose/PBS solution. Flies were fed on PQ on alternating days, for 8 days in total (final age = 22–25 days), prior to dissection and imaging. Flies were returned to standard food on days without drug treatment to minimise the impact of a sucrose-only diet.

Flies fed with 5 mM PQ had significantly increased the abundance of MT unbundling within T1 axons, which correlated with an increase in axonal swellings (**Figure S1**), confirming the detrimental effect of increased ROS on MTs and axonal health. However, in PQ-treated flies, overexpression of catalase fully rescued PQ-induced MT-associated phenotypes (**Figure S1**). Despite the protective effect observed from catalase overexpression on MTs when exposed to PQ, it was not able to change the number of axonal swellings (**Figure S1**).

Our second approach aimed to enhance the cellular antioxidant defence system by expressing *UAS-Keap1 RNAi to* increase NRF2 activity (Stapper and Jahn, 2018). We found that *Keap1* knockdown in T1 neurons substantially reduced the frequency of MT unbundling and axonal swellings (**Figure 4**). In addition, Keap1 knockdown halted the morphological deterioration of synaptic terminals, with a significantly greater number of MTs observed at synaptic sites compared with age-matched control flies (**Figure 4**). Taken together, these findings indicate knockdown of Keap1 prevents age-related MT deterioration and correlates with a reduction in axonal and synaptic atrophy. However, upregulating catalase only provides some beneficial effects to MT integrity in conditions of elevated ROS but has an overall detrimental effect on the health of axons.

**Figure 4.**
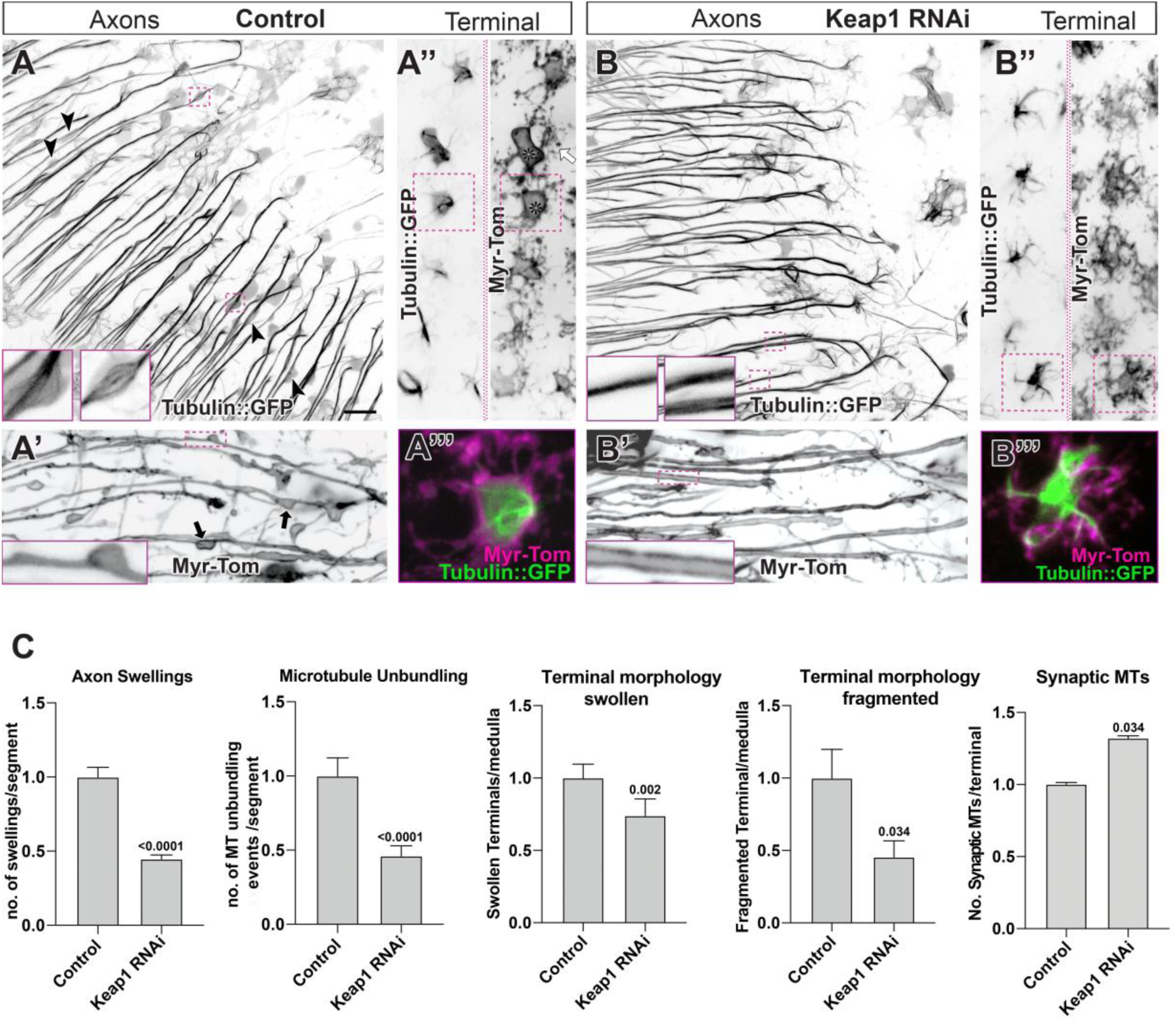
Knocking down of Keap1 attenuates the onset of age-related microtubule alterations and morphological changes in axons and terminal. (A–B’) Medulla regions of adult brains aged 30-40 days post eclosure, which show T1 axons labelled with GFP-tagged α-tubulin (tubulin::GFP) and the plasma membrane marker myristoylated-Tomato (myr-Tom). Aged neurons in the absence (A) or presence of Keap1 knockdown (*Keap1 RNAi*; B) are compared. In axons (A–B’) *Keap1* knockdown supresses the appearance of microtubule unbundling and axonal swellings (black arrow heads for microtubule unbundling and black arrows for swellings in A and A’ compared with B and B’). *Keap1* knockdown also improves axonal terminals (A’’-B’’’) which contains more synaptic microtubules and less swollen and broken terminals (asterisk for swollen and white arrows for broken synapses in A’’ compared to B’’ and insets with magnified images of a terminal in A’’’ and B’’’). (**C**) Quantifications of phenotypes described above; bars represent normalised mean ± SEM; p values are shown above each bar, as assessed by Mann-Whitney rank sum tests. For the assessment of axon swellings and microtubule unbundling, data were collated across two individual repeats, from a minimum of 300 axonal segments from 15 control and 20 Keap1 RNAi medullas. For the assessment of axon terminal phenotypes, data were collated across four individual repeats, from a minimum of 700 synapses from 41 control and 47 Keap1 RNAi medullas. Scale bars = 10 μm.

### Oxidative stress alters microtubule organisation and stability in primary neuronal cultures

Given the changes in MTs observed by our *in vivo* ageing experimental conditions of increased ROS, we proposed that MT-based physiology may be a key target of oxidative stress. To determine the processes whereby elevations in ROS impact MT physiology, we used *Drosophila* primary neuronal cell cultures, established in previous work, as an *in vitro* neuronal model (Sánchez-Soriano et al., 2010).

PQ, which promotes mitochondrial superoxide radicals, and diethyl maleate (DEM), which depletes GSH levels in cells (Cochemé and Murphy, 2008, Mitchell et al., 1983) have been previously utilised in research on oxidative stress (Dasgupta et al., 2012, Albano et al., 2015). Here, we cultured *Drosophila* embryo derived primary neurons, which were treated with either vehicle, 100 μM PQ or 100 μM DEM at 3 days in vitro (DIV) and fixed and imaged at 6 DIV. A MT disorganisation index (areas with MT disorganisation/axon length) was employed as a robust readout indicative of alterations in MT organisation (Hahn et al., 2021, Qu et al., 2017). In both PQ- and DEM-treated neurons, prominent MT curling and unbundling within axons was observed in comparison with vehicle-treated neurons (**Figure 5**). This ‘MT disorganisation’ phenotype shared strong similarity to MT abnormalities observed in ageing T1 neurons (**Figure 1**). To confirm whether this physiological disruption to MTs was mediated by excess ROS formation, we also co-treated neurons with the vitamin E analogue, Trolox, with PQ or DEM. Trolox is a potent ROS scavenger that exerts antioxidant effects (Brigelius-Flohé and Traber, 1999, Giordano et al., 2020). Here, co-administration of Trolox fully prevented the development of PQ- and DEM-induced MT disorganisation, indicating that dysregulation of MT organisation was mediated by elevated levels of ROS (**Figure 5**).

**Figure 5.**
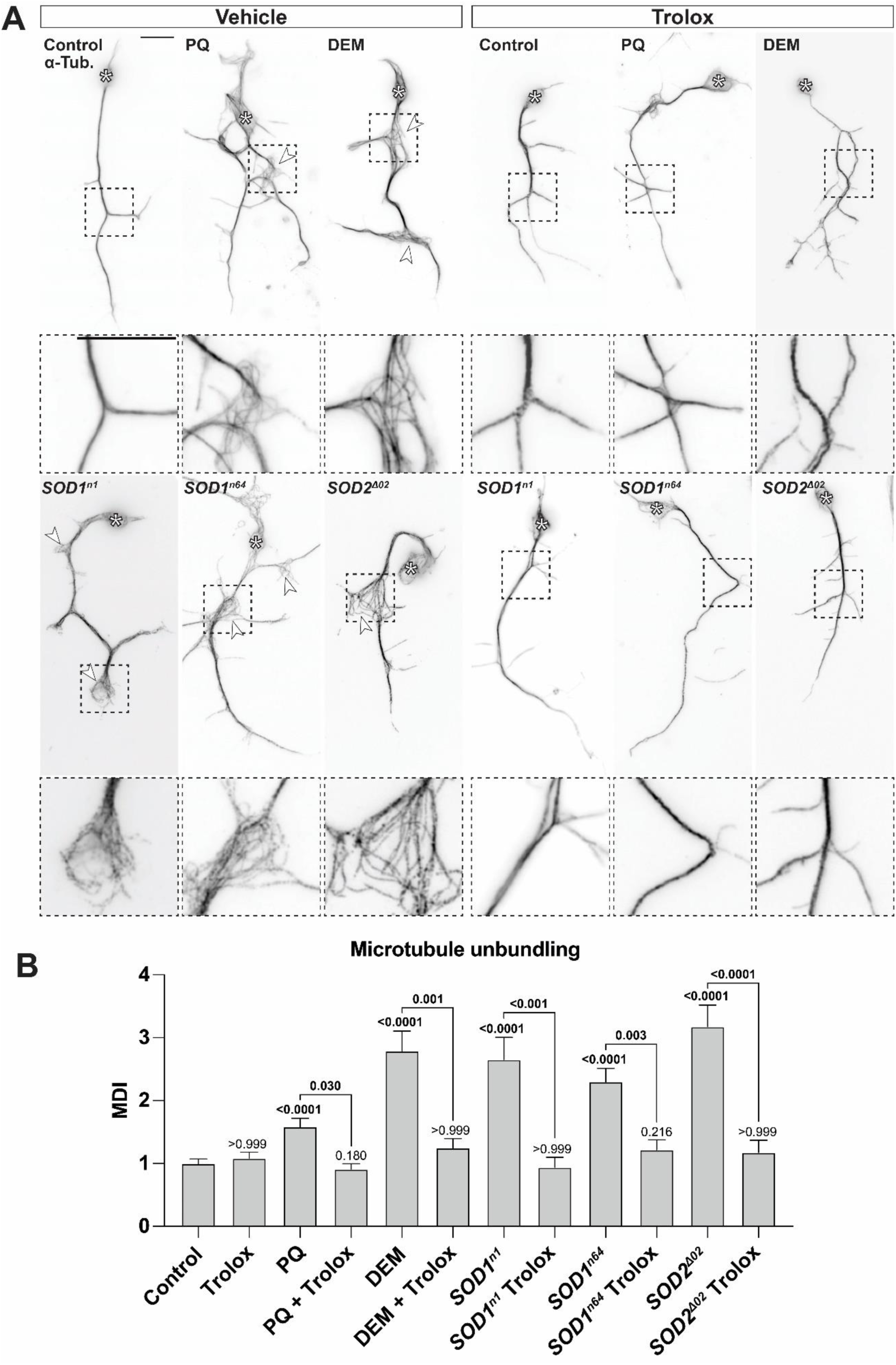
ROS promote microtubule disorganisation in primary neuronal cultures. (A) Representative images of primary *Drosophila* primary neurons 6 DIV stained for tubulin. Neurons at different conditions: control treated with 100 μM PQ, control treated with 100 μM DEM, and neurons carrying either of the mutant alleles *SOD1^n1^*, or *SOD1^n64^* or *SOD2^Δ02^* in homozygous. Cells were treated with ethanol (vehicle) or 100 μM trolox. White arrow heads indicate regions of microtubule disorganisation, magnified in dashed boxes. (B) Quantitative analysis of the microtubule disorganisation index (MDI). Bars represent normalised mean ± SEM. P values are shown above each bar, as assessed by a Kruskal-Wallis one-way test. Data were collated across 14 experiments with 3 individual cultures per condition per repeat, with a minimum of 100 overall neurons evaluated per condition. Scale bars = 10 μm.

To further validate these findings and discard confounding toxicity or off-target effects from PQ and DEM as a potential cause of these phenotypes, we next tested whether neurons deficient in SOD activity displayed similar MT aberrations. Despite lethality in the adult, harvesting neurons from *Drosophila* embryos facilitated the culturing of cells harbouring two-copies of mutations within SOD genes. Two amorphic mutant SOD1 alleles (*SOD1^n64^* and *SOD1^n1^;* (Phillips et al., 1995)) were tested. Neurons harbouring either mutation exhibited severe MT curling and unbundling (**Figure 5**). Likewise, we found that homozygous *SOD2^Δ02^* neurons had significantly increased MT disorganisation, suggesting that loss of SOD antioxidant activity within the cytosol or mitochondria was sufficient to disrupt MT bundling *in vitro*. Trolox supplementation, administered during plating (0 DIV) and refreshed at 3 DIV, significantly prevented the onset of this MT disorganisation in both SOD1- and SOD2-mutant neurons (**Figure 5**).

Given the dramatic change in MT organisation in response to oxidative stress, we investigated whether ROS impacts MT stability using a previously reported nocodazole-sensitivity assay in primary *Drosophila* neurons as a proxy readout (Szikora et al., 2017, Voelzmann et al., 2016). Nocodazole is a reversible agent that through its interaction with β-tubulin inhibits MT assembly (Szikora et al., 2017). Since MT polymerisation is inhibited in the presence of nocodazole, it is possible to assess whether existing MTs are labile or stable.

In primary neurons incubated with 100 μM nocodazole for 5.5 hours, the number of regions within axons where MTs were absent (as consequence of depolymerisation) were recorded and quantified per cell. These regions devoid of MTs often appear as ‘gaps’ or ‘breaks’ within MT bundles and were used to infer changes in MT stability or resistance to nocodazole. MT gaps could be seen in control cells treated with nocodazole. Significantly greater numbers of MT gaps were observed in SOD1- or SOD2-mutant neurons, suggesting that excessive ROS renders MTs less stable (**Figure S2**). Additionally, we found a small exacerbation in the number of MT gaps in neurons co-treated with nocodazole and DEM for 5– 6 hours. However, in neurons pre-treated with DEM for a further 12–13 hours, prior to nocodazole treatment (still co-treated with DEM), a substantial increase in the number of MT gaps was observed compared with nocodazole alone. This elevation in MT gaps was significantly enhanced in comparison with the shorter DEM incubation time (**Figure S2 A and C**).

Taken together, these data suggest that excessive ROS disrupts the organisation and decreases the stability of axonal MTs.

### EB1 mediates MT alterations induced by oxidative stress in cultured neurons

At this stage, we had established a clear relationship between increasing levels of ROS and disrupted MT physiology; however, the mechanism(s) whereby ROS impacts MTs is unknown.

MT polymerisation predominantly occurs at the MT plus end, and is regulated by the family of MT plus end-binding factors (EBs) (Goodson and Jonasson, 2018). The high affinity of EBs for GTP-bound tubulin results in their transient binding to the plus ends of growing MTs, forming dynamic and characteristic ‘comet’-like foci. *In vitro* polymerisation assays have demonstrated that EB1 regulates MT dynamics and stability, and investigations in *Drosophila* neurons showed that a reduction in EB1 comet size correlates with an increase in MT disorganisation (Duellberg et al., 2016, Hahn et al., 2021, Rickman et al., 2017). Given EB1 is a key MT regulator in *Drosophila* neurons, we hypothesised that the effects of oxidative stress observed on MTs could be mediated through changes in EB1.

To investigate the impact of ROS on EB1 function, we first assessed the ability of EB1 to bind to MT plus ends. We visualised endogenous EB1 comets in primary neuronal cultures by immunohistochemistry and measured the mean comet length as a proxy measure for EB1 recruitment to MT plus ends. Control and SOD1 or SOD2 mutated neurons were compared at 6 hours in vitro, because at this timepoint, EB1 comets are prominent and easy to visualise. We found that in neurons homozygous for *SOD1^n64^*, *SOD1^n1^* or *SOD2^Δ02^*, axonal EB1 comet length was significantly reduced compared with control neurons (**Figure 6**). Furthermore, comet length was fully rescued when supplemented with Trolox at plating (**Figure 6**), suggesting that increasing ROS levels are inversely associated with EB1 comet size. We therefore proposed that ROS impair EB1 recruitment on MTs and hypothesised that MT defects induced by oxidative stress are driven by EB1 changes at the MT plus end.

**Figure 6.**
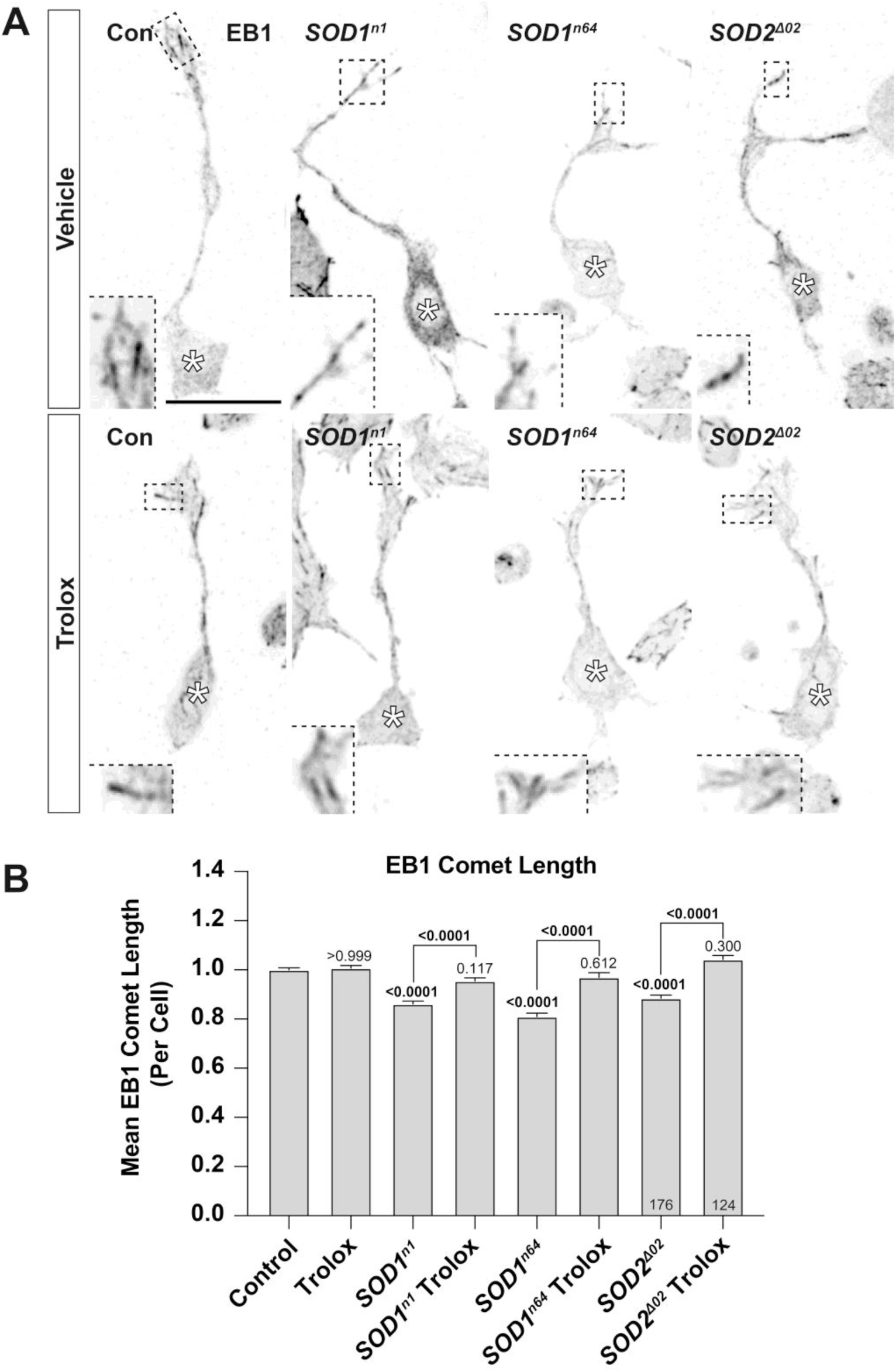
ROS reduces EB1 comet length in primary neurons. (A) Representative images of primary *Drosophila* primary neurons 6 HIV stained for EB1. Neurons of different genotypes: *w^1118^* (control) or carrying the *SOD1^n1^* or *SOD1^n64^* or *SOD2^Δ02^* mutant allele in homozygous. Cells were treated with ethanol (vehicle) or 100 μM trolox during plating (0 HIV). Asterisks indicate cell bodies and dashed boxes show regions of EB1 comets, magnified in the respective inset. (B) Quantitative analysis of EB1 comet lengths per cell; bars represent mean ± SEM normalised to control; P values are shown above each bar, as assessed by a Kruskal-Wallis one-way test. Data were collated across six independent repeats with 3 individual cultures per condition per repeat, with a minimum of 120 neuros evaluated per condition. Scale bar = 10 μm.

To confirm our hypothesis, we considered that enhancing EB1 activity or affinity for the MT plus end will rescue ROS-induced reductions in comet size, and in-turn ROS-induced MT defects. To test this, we utilised the neuroprotective peptide NAPVSIPQ (NAP), derived from activity-dependent neuroprotective protein (ADNP), which exhibits an EB1-interacting SxIP motif (Oz et al., 2012, Oz et al., 2014). Previously it was shown in differentiated neuroblastoma N1E-115 cells that NAP strongly enhances EB dynamics and comet length, through modulation of EBs function (Ivashko-Pachima et al., 2017). In SOD1- and SOD2-mutant neurons, we further found that 1 nM NAP treatment fully rescued ROS-induced EB1 comet shortening compared with mutant control neurons (**Figure 7**). Remarkably, NAP supplementation also fully protected neurons against ROS-induced MT disorganisation (**Figure 8**). These data suggest that dysregulation of MTs by oxidative stress can be rescued by supporting EB1 binding at plus ends in the absence of boosted antioxidant capacity, and therefore changes in EB1 function may be responsible for ROS-induced MT unbundling. Supportive of these findings, we also found that overexpressing EB1 (*elavGal4; UAS-Eb1::GFP*) in primary neurons rescued DEM-induced MT disorganisation (**Figure S3**).

**Figure 7.**
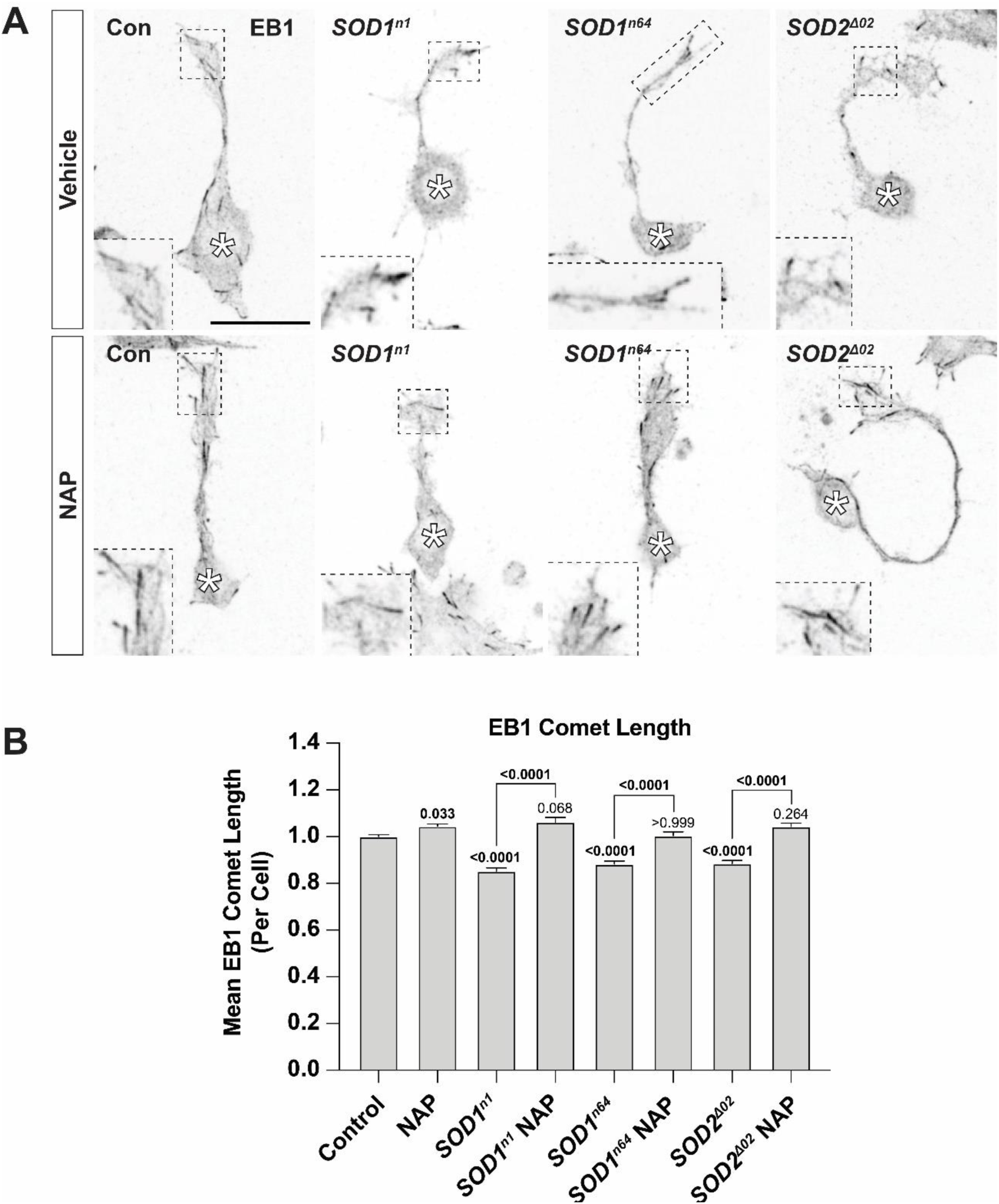
NAP rescues ROS-induced shortening of EB1 comet length. (A) Representative images of primary *Drosophila* neurons cultured from *w^1118^* (Control), homozygous *SOD1^n1^*-, *SOD^1n64^-* or *SOD2*^Δ*02*^*-*mutant embryos, stained for EB1 at 6 HIV. Cells were treated with H_2_O (vehicle) or 1 nM NAP during plating (0 HIV); asterisks indicate cells bodies and dashed boxes show regions of EB1 comets, magnified in the respective inset. (B) Quantitative analysis of EB1 comet lengths per cell; bars represent normalised mean ± SEM. P values are shown above each bar, as assessed by a Kruskal-Wallis one-way test. Data were collated across six independent repeats with 3 individual cultures per condition per repeat, with a minimum of 60 neurons evaluated per condition. Scale bar = 10 μm.

**Figure 8.**
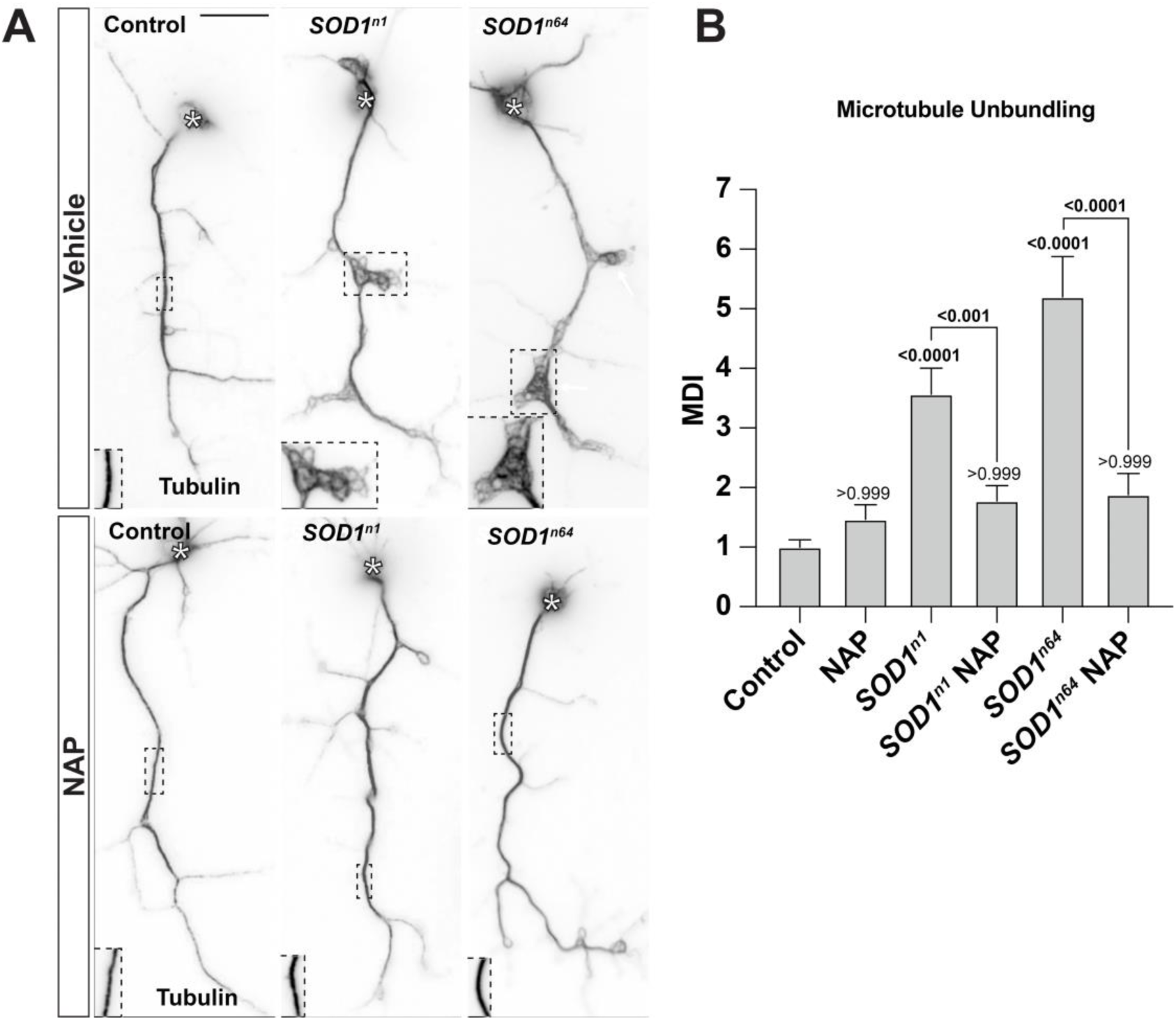
NAP rescues ROS-induced microtubule disorganisation in primary neurons. (A) Representative images of primary *Drosophila* neurons stained for tubulin. Neurons of different genotypes: *w^1118^* (control) or carrying the *SOD1^n1^*- or *SOD1^n64^*-mutant allele in homozygous. Cells were treated with H_2_O or 1 nM NAP at 6 HIV and refreshed at 3 DIV; asterisks indicate cells bodies, regions of microtubule disorganisation within dashed boxes are shown magnified in the respective inset. (B) Quantitative analysis of MDI per cell. Bars represent normalised mean ± SEM. P values are shown above each bar, as assessed by a Kruskal-Wallis one-way test. Data were collated across four independent repeats with 3 individual cultures per condition per repeat and over 120 neurons per condition. Scale bar = 10 μm.

### EB1 protects against oxidative stress *in vivo*

Given that our data generated in cultured neurons show that enhancing EB1 protects against the effects of ROS, we lastly tested whether the same relationship could be observed *in vivo* using the T1 ageing model. We utilised the protocol described above to feed PQ and DEM to flies and evaluate its effects in the T1 model. Both 2.5 mM DEM or 5 mM PQ treatment resulted in significant enhancement of ageing phenotypes in T1 cells, including MT unbundling, axonal swellings, and thinning, and swollen synaptic terminals (**Figure 9** and **Figure 10**). We then challenged flies overexpressing EB1 (*UAS-EB1::mCherry*) in T1 neurons with 2.5 mM DEM or 5 mM PQ, and strikingly found that EB1 overexpression fully suppressed PQ and DEM induction of MT unbundling and markedly reduced the frequency of axonal swellings (**Figure 9**). In addition to this, we also observed widespread improvements in synaptic morphology and synaptic MTs. Overexpression of EB1 partially rescued the number of invading synaptic MTs and proportion of terminals with broken MTs compared with non-EB1 overexpression controls (**Figure 10**). EB1 overexpression also rescued PQ-induced synaptic terminal swelling (**Figure 10**).

**Figure 9.**
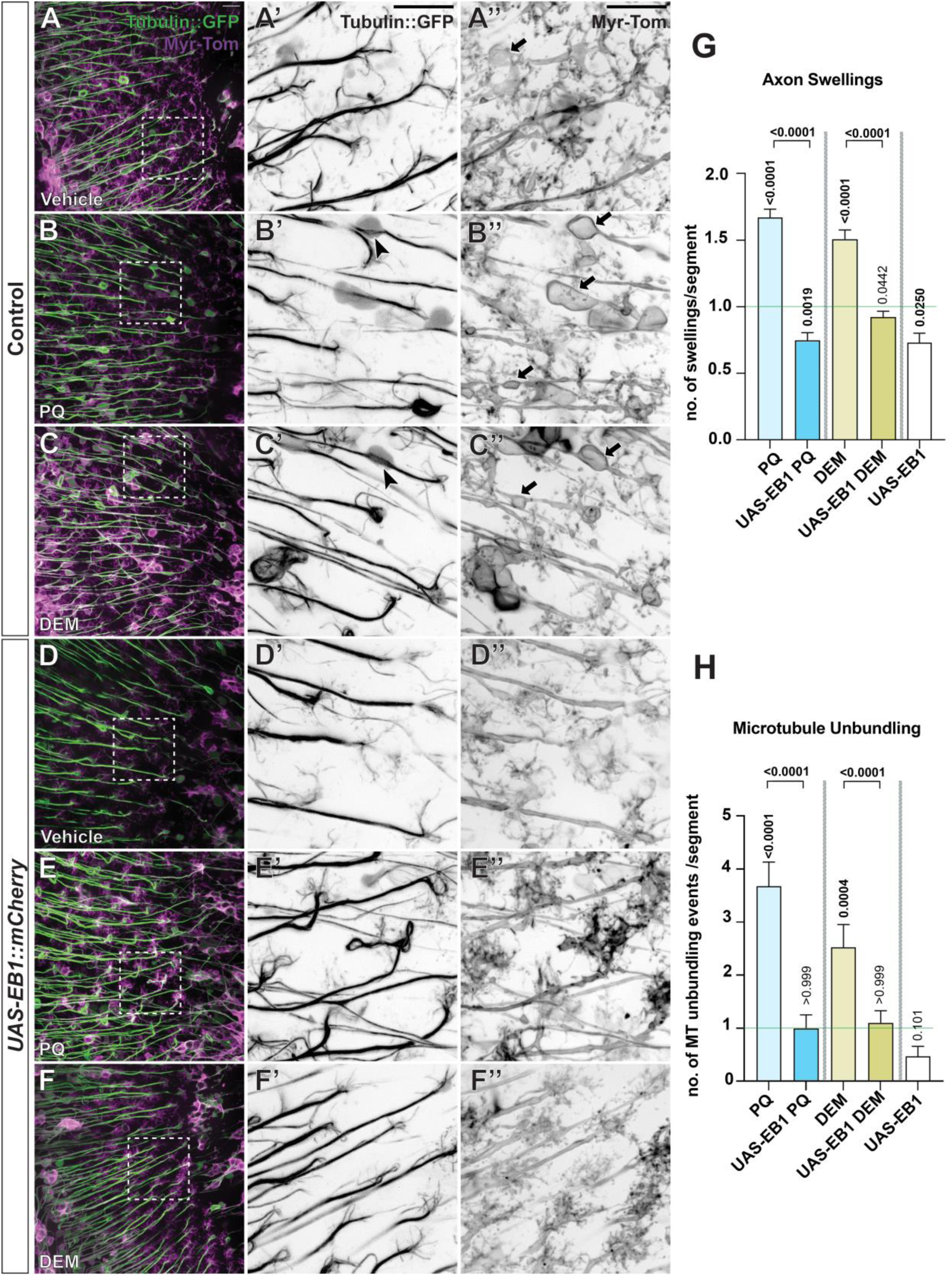
Overexpression of EB1 prevents oxidative stress-induced microtubule unbundling and axonal swellings within the aged brain. (A–F’’) Medulla region of adult brains at 22–25 days post eclosure, depicting T1 axons labelled with GFP-tagged α-tubulin (tubulin::GFP) and the plasma membrane marker myristoylated-Tomato (myr-Tom). At 14–17 days post eclosure, flies were treated with vehicle (A, D) or 5 mM PQ (B, E) or 2.5 mM DEM (C, F) in 2.5% sucrose every alternate day. Magnified images of regions, outlined by dashed white boxes are shown for tubulin::GFP (A’, B’, C’, D’, E’, F’) and Myr-Tom (A’’, B’’, C’’, D’’, E’’, F’’) as inverted greyscale images. Aged neurons in the absence (A–C’’) or presence (D–F’’) of EB1::mCherry overexpression are compared. *EB1* overexpression supresses oxidative stress induced phenotypes in ageing neurons, comprising axon swellings (arrows) and MT unbundling (arrow heads); boxed areas shown as magnified images below. Black arrows indicate regions of axonal swellings and black arrow heads indicate regions of microtubule disorganisation. (G–H) Quantitative analyses of the frequency of axonal swellings and microtubule unbundling. Conditions with drug treatment have been normalised and compared to internal vehicle-treated controls (green line). Bars represent normalised mean ± SEM; P values are shown above each bar, as assessed by Kruskal-Wallis one-way tests. Data were collated across 4 repeats for experiments involving PQ and 5 repeats in experiments involving DEM. Number of medullas assessed per group were 35 in PQ (5 mM), 32 in DEM (2.5 mM), 24 in UAS-EB1::mCherry, 21 in UAS-EB1::mCherry PQ and 38 in UAS-EB1::mCherry DEM. A minimum of 300 axonal segments were evaluated per condition. Scale bars = 10 μm.

**Figure 10.**
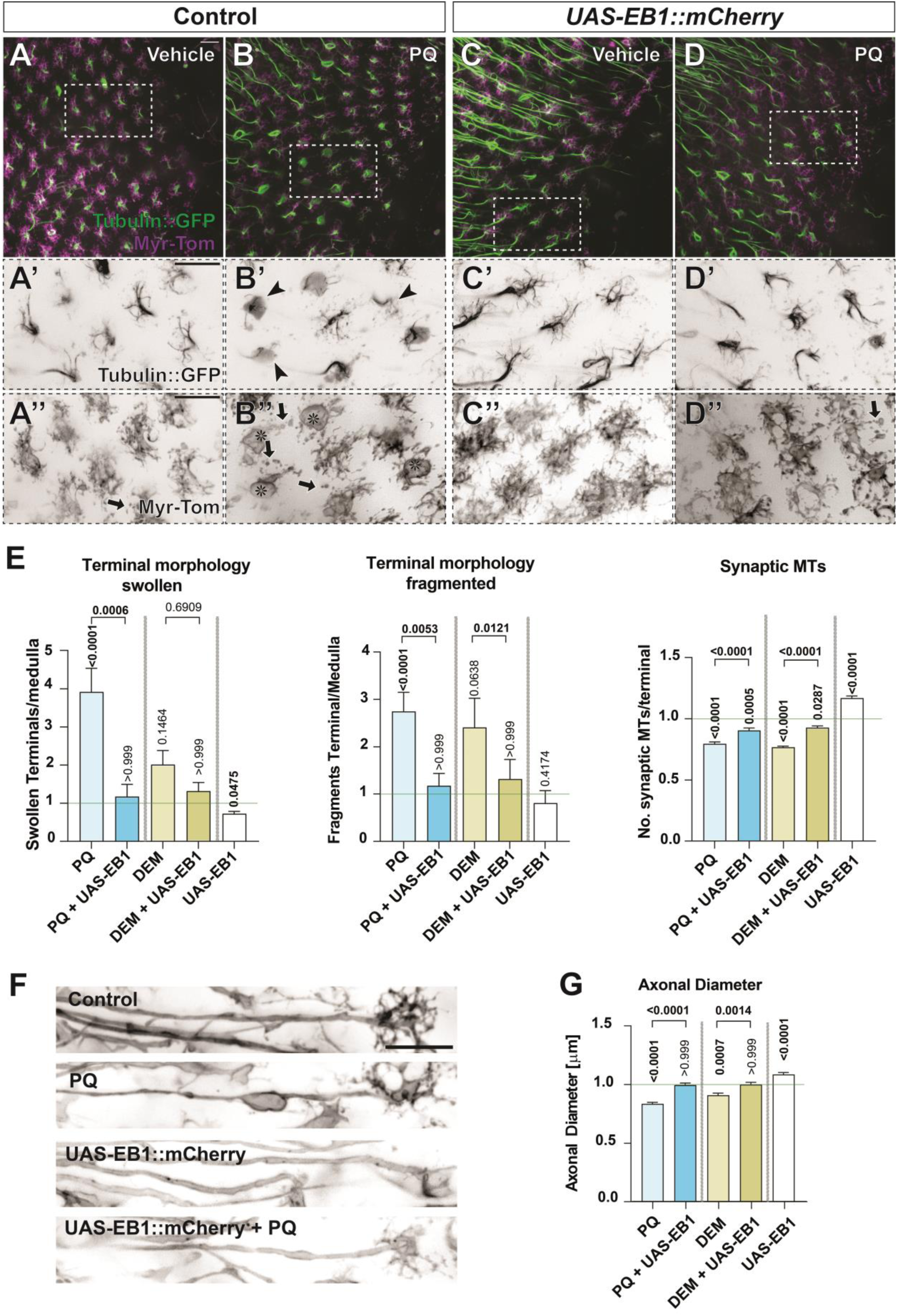
Oxidative stress-induced synaptic terminal aberrations and axonal thinning is prevented by overexpression of EB1. ((A–D) Medulla region of adult brains at 22–25 days post eclosure, depicting T1 axons labelled with GFP-tagged α-tubulin (tubulin::GFP) and the plasma membrane marker myristoylated-Tomato (myr-Tom). At 14–17 days post eclosure, flies were treated with vehicle (A, C) or 5 mM PQ (B, D) in 2.5% sucrose every alternate day. Magnified images of regions, outlined by dashed white boxes are shown for tubulin::GFP (A’, B’, C’, D’) and Myr-Tom (A’’, B’’, C’’, D’’) as inverted greyscale images for easier visualisation. Aged neurons in the absence (A–B’’) or presence (C–D’’) of EB1::mCherry overexpression and with different oxidant treatments are compared. PQ induced ageing-related phenotypes at the synaptic terminals, including an increase in swollen (asterisks) and broken terminals (arrows), and a decrease in synaptic MTs (arrowheads), are supressed upon *EB1* overexpression in T1 neurons. (F) Representative images of T1 axonal membranes labelled with myristoylated-Tomato and under same conditions as indicated above. Note PQ induces a decrease of the diameter of axons which can be counteracted by EB1::mCherry overexpression. (E and G) Quantitative analyses using the conditions above as well as DEM treatments, of the proportion of synaptic terminal swellings per medulla, fragmented synaptic terminals per medulla, mean number of synaptic microtubules per terminal and axonal diameter. Conditions with drug treatment have been normalised and compared to internal vehicle-treated controls (green line). Bars represent normalised mean ± SEM; P values are shown above each bar, as assessed by Kruskal-Wallis one-way tests. Data were collated across 4 individual repeats per condition, with a minimum of 20 medullas evaluated per condition. Scale bars = 10 μm.

Previous work has shown that axon diameter is impacted by ageing, and is also evident in the T1 ageing model (Okenve-Ramos et al., 2024). Consistent with PQ and DEM exacerbating ageing hallmarks, we observed a significant decrease in mean axonal diameter compared with age-matched, vehicle-fed control flies (**Figure 10**). Overexpression of EB1 prevented PQ-and DEM reductions in axonal diameter (**Figure 10**).

In summary, these data show that EB1 overexpression protects against oxidative stress *in vivo*. Furthermore, that axonal and synaptic deterioration induced by oxidative stress is caused by the dysregulation of the MT cytoskeleton, with which MT plus end physiology is significantly implicated.

## Discussion

Here, we aimed to investigate mechanisms whereby oxidative stress may impact neuronal physiology and ageing. Our results showed that both pharmacological (PQ and DEM) and genetic-driven (SOD2 mutant strain; SOD1 and GSS1 and 2 knockdown) induction of oxidative stress exacerbates hallmarks of ageing in T1 neurons, which included the onset of axonal swellings, reduction in axonal diameter, and morphological transformation of axonal synaptic terminals. The discovery that MT deterioration observed in the natural course of the ageing brain (Okenve-Ramos et al., 2024), is exacerbated by oxidative stress, led to a focus on the interplay between ROS and MT regulation. We found that oxidative stress led to a prominent reduction in the binding of EB1 at MT plus ends. By increasing EB1 activity, through the addition of NAP or EB1 overexpression, ROS-induced MT disorganisation could be fully prevented *in vitro*. A remarkable finding was that *in vivo* overexpression of EB1 using the ageing T1 cell model fully prevented PQ-induced ageing hallmarks in both axonal and synaptic compartments.

### Phenotypic hallmarks of neuronal ageing are increased by oxidative stress

The onset of focal axonal swellings is considered a classical hallmark of aged nerve tissue and an early marker of Wallerian degeneration (Coleman, 2005). Prominent neuronal swellings, or blebbing, has been observed in aged brain tissue from mice, rat, primate, and humans (Takeuchi et al., 1995, Schmidt et al., 1997, Fiala et al., 2007, Marangoni et al., 2014). Notably, these swellings observed here in the *Drosophila* optic lobe resemble such cases found in other higher order animal models, suggesting convergent pathological consequences across species. High rates of axonal swellings are also observed in many neurodegenerative conditions including AD, PD, ALS and multiple sclerosis as well as in cases of traumatic brain injury (TBI) (Cheng and Povlishock, 1988, Trapp et al., 1998, Galvin et al., 1999, Tsai et al., 2004, Denton et al., 2016), with axon swellings being a typical sign of axonopathy (Marangoni et al., 2014, Prokop, 2021), hence its value as a readout for disease severity and onset of age-related neurodegeneration. Our data show that axonal swellings during ageing are promoted by increasing ROS levels. This was supported through the depletion of SOD activity and GSH production, which accelerated ageing phenotypes. Additionally, by reducing Keap1 levels, thus enhancing NRF2 activity, age-related axonal swellings were reduced.

Interestingly, PQ, as used here, is a well-known herbicide and neurotoxicant which is associated with the onset of sporadic PD (Vaccari et al., 2017, Vaccari et al., 2019, Weed, 2021). The effects observed here in the context of ageing may substantiate the impact of ROS in neurodegenerative disorders, such as PD. Further to this, in patients with traumatic brain injury (TBI), nitrogen oxide (NOX) activity is elevated as early as one-hour post-injury, suggesting a pathological role of ROS in the outcomes of TBI (Zhang et al., 2012); the extent of ROS-associated tissue damage and mitochondrial dysfunction in TBI patients correlates with more severe outcomes (Tavazzi et al., 2005, Valko et al., 2007). This increase in ROS following brain injury may trigger degenerative signalling pathways, predisposing individuals to AD and other neurodegenerative conditions (Breunig et al., 2013, Klomparens and Ding, 2020, Mendez, 2017).

### Oxidative stress, microtubule dysregulation and axonal atrophy

Another key finding here was the onset of MT unbundling by inducing ROS, which is an early ageing hallmark observed in the T1 model (Okenve-Ramos et al., 2024). In aged rhesus monkey brains, unbundled and disorganised MTs have also been observed in axonal swellings, patently reminiscent of MT unbundling in T1 cells (Fiala et al., 2007). MT reduction has also been observed in AD as well as ageing controls as assessed in brain biopsies (Cash et al., 2003), with this MT reduction being independent of Tau pathology. Given these observations across organisms, we propose an ageing cascade whereby oxidative stress triggers MT deterioration, which in turns drives axonal atrophy. Several lines of research demonstrate that MTs are vital for axonal integrity. Our past work has established that reducing the function of several MT binding proteins led to higher rates of axonal swellings during ageing, and inversely, enhancing MTs decreased the prevalence of axonal swellings during ageing (Okenve-Ramos et al., 2024).

The causative connection between MT deterioration and axonal swellings is not a specific trait of insect neurons, as this physiology has also been observed in vertebrates. For instance, nocodazole-induced MT disassembly in chick dorsal root ganglia (Datar et al., 2019), or MT decay caused by loss of MAP1b in mouse Purkinje neurons (Liu et al., 2015), have both been shown to promote the formation of axonal swellings. The ability of ROS to modify MT networks may be a common phenomenon found in higher organisms and in different cell types; for example, disorganised MTs are evident in human osteosarcoma cells and in human umbilical cord vein endothelial cell cultures treated with H_2_O_2_ (Lee et al., 2005, Valen et al., 1999). Further, elevated ROS in myocytes have been shown to promote MT depolymerisation (Drum et al., 2016).

Our work suggests that ROS affects MT regulation through a mechanism whereby excessive ROS hinders the ability of EB1 to bind to MT plus ends. We find that ROS-induced deficits in EB1 activity and consequently in MT organisation can be supressed by increasing EB1 levels or by supplementing neurons with the small peptide, NAP. NAP is derived from ADNP (Amram et al., 2016, Bassan et al., 1999), and previous work has shown that NAP rescues axonal transport and MT deficits in a *Drosophila* tauopathy model (Quraishe et al., 2013). Furthermore, in a SOD1-G93A mouse model of ALS, NAP protected against disrupted axonal transport (Jouroukhin et al., 2013), and in PC12 cells, NAP has also demonstrated protective effects against oxidative stress (Steingart et al., 2000). Our current study provides a possible mechanistical explanation as to how NAP protects against elevated ROS based on our observations on how ROS and NAP affect EB1 and MTs in neurons. NAP (investigational drug: davunetide; also known as CP201) is under clinical investigation for neurodegenerative, MT-related tauopathies (Gozes et al., 2023) as well as the autism-related ADNP syndrome (Gozes, 2020).

The mechanisms of EB-dependent ROS-induced damage could also extend to other cell types. For instance, in A549 cells and ventricular myocytes, ROS reduced accumulation of EB1 at MT plus ends (Drum et al., 2016, Le Grand et al., 2014). How ROS mediates its effects on MT-binding factors is open for discussion; potential mechanisms could include direct thiol oxidation (Landino et al., 2007, Landino et al., 2004) or activation of redox-sensitive signalling pathways that are involved in the regulation of MT, such as c-Jun N-terminal Kinase (JNK), apoptosis signal-regulating kinase (ASK1) (Daire et al., 2009), glycogen synthase kinase 3 (GSK3) (Hur et al., 2011) and p38 mitogen-activated protein kinases (p38 MAPK).

### The origin of ROS production in ageing

One of the challenges in understanding the role of oxidative stress is ageing is determining the source of ROS which is implicated in ageing. For example, studies have shown that ROS elevations can enhance lifespan (Ristow and Schmeisser, 2011), which contradicts research of the classical theories of oxidative stress in ageing. Our data using the T1 ageing model suggest that SOD2 heterozygote loss*-*of*-*function exhibits a greater acceleration of ageing hallmarks compared with SOD1 mutant heterozygotes; although, a mild increase in axonal swellings from the knockdown of SOD1 was observed. This suggests that mitochondrial-localised ROS may have a greater impact on the ageing process in the brain. Supporting this, is the finding that loss of Milton/Miro-mediated mitochondrial transport enhances MT disorganisation in primary neuronal cultures (Liew et al., 2021). In addition, targeted knockdown of Gss1 and 2 in T1 neurons also accelerated ageing phenotypes, hence loss of glutathione metabolism could be implicated in the ageing process, and critically, that these effects are cell autonomous. A recent study using Drosophila primary neuronal cultures confirmed that, in-addition to SOD1 and SOD2 mutants, catalase loss-of-function and overexpression of NOX or Duox also promotes MT unbundling in cultures neurons (Liew et al., 2021). Given these data, this could suggest that ROS-induced MT disorganisation may not necessarily be driven by subcellular specific origins of ROS.

We found that depletion of GSH via DEM treatment also strongly induced MT unbundling and decreased MT stability. Interestingly, previous research has shown that the depletion of mitochondrial GSH pools in cerebellar granule neurons (CGNs) stimulates ROS production and apoptotic pathways (Wilkins et al., 2013). Moreover, CGNs (and astrocytes) are sensitive to GSH depletion in mitochondrial pools, with less of an impact observed when depleting only cytosolic GSH, suggesting that mitochondrial GSH content may be more fundamental for redox homeostasis, and targeted by DEM (Muyderman et al., 2007, Wüllner et al., 1999).

Further supporting the importance of GSH in healthy ageing, we found that DEM exacerbated axonal swellings and other ageing hallmarks in the T1 ageing model. Reduced levels of GSH are associated with apoptosis in degenerating neurons (Diaz-Hernandez et al., 2005, Merad-Boudia et al., 1998). Furthermore, GSH depletion and elevated ROS is suspected to be an early preclinical marker of neuronal death in the substantia nigra during PD onset (Dexter et al., 1994, Merad-Boudia et al., 1998), and altered GSH metabolism is apparent in hippocampal and cerebellar tissue from patients with AD (Aksenov and Markesbery, 2001).

In summary, the findings described here support a role of oxidative stress as a key driver of neuronal ageing and suggest a novel role of ROS in sensitising the MT plus end. Manipulating MT-associated factors such as EB1 could be a novel and valuable therapeutic strategy to prevent the neurological decline observed with age, and that EB1 could also be a relevant therapeutic target in disease conditions with intrinsic oxidative stress.

## Acknowledgments and disclosures

This work was made possible through support to N.S.S by the BBSRC (BB/R018960/1, BB/W016907/1), to S.S and N.S.S by the Wellcome Trust (WT204002). We thank the University of Liverpool Imaging Facility for assistance during the project. We also thank Hiro Ohkura for kindly providing the DmEB1 antibody. Stocks obtained from the Bloomington *Drosophila* Stock Center (NIH P40OD018537) were used in this study. Davunetide was provided by Illana Gozes and is currently under patent protection and clinical development by ExoNavis Therapeutics Ltd. I.G. VP Drug Development.

