## Supplementary figure 1 for "Oxidative stress promotes axonal atrophy through alterations in microtubules and EB1 function"

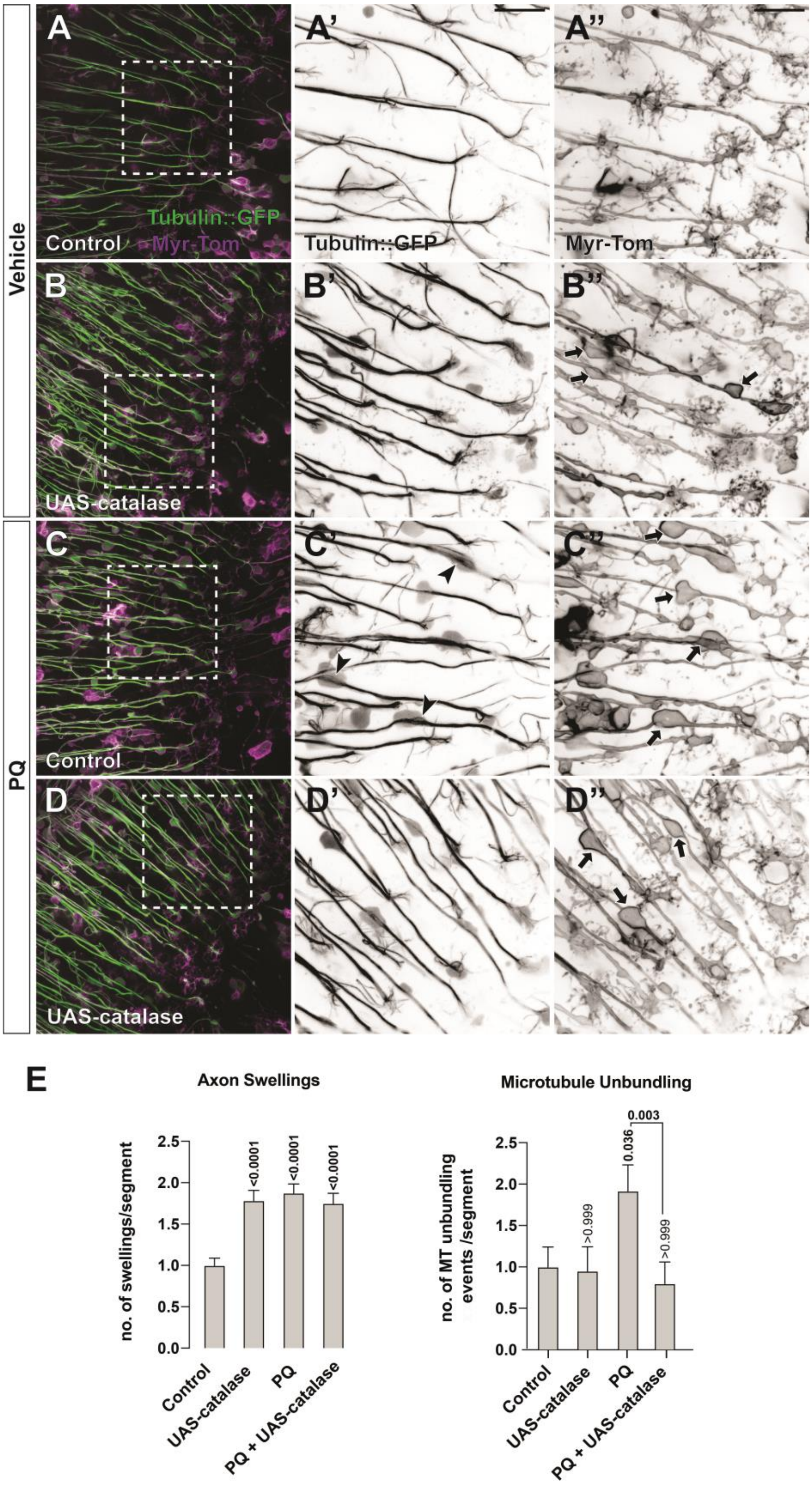


**Figure S1. Catalase overexpression rescues PQ microtubule disorganisation**
(A–D'') Medulla region of adult brains at 22–25 days post eclosure, depicting T1 axons labelled with GFP-tagged α-tubulin (tubulin::GFP) and the plasma membrane marker myristoylated-Tomato (myr-Tom). Flies without or with UAS-catalase expression in T1 neurons were treated at 14–17 days post eclosure with $H_2O$ (A–B'') or 5 mM PQ (C–D'') in 2.5% sucrose every alternate day. Magnified images

of regions, outlined by dashed white boxes are shown for tubulin::GFP (A', B', C' and D') and Myr-Tom (A'', B'', C'' and D'') as inverted greyscale images for easier visualisation. PQ induces microtubule disorganisation in axons which is prevented by catalase overexpression (black arrow heads in C' compared to D'). PQ also induces axonal swellings which cannot be rescued by catalase overexpression (black arrow in C'' and in D''). (E) Quantitative analyses of the phenotypes above. Bars represent normalised mean ± SEM; P values are shown above each bar, as assessed by Kruskal-Wallis one-way tests. Two individual repeats were performed with a total of 18 control, 12 UAS-catalase, 22 5 mM PQ and 17 UAS-catalase + PQ medullas assessed. A minimum of 180 axonal segments were evaluated. Scale bars = 10 µm.
