## Supplementary figure 2 for "Oxidative stress promotes axonal atrophy through alterations in microtubules and EB1 function"

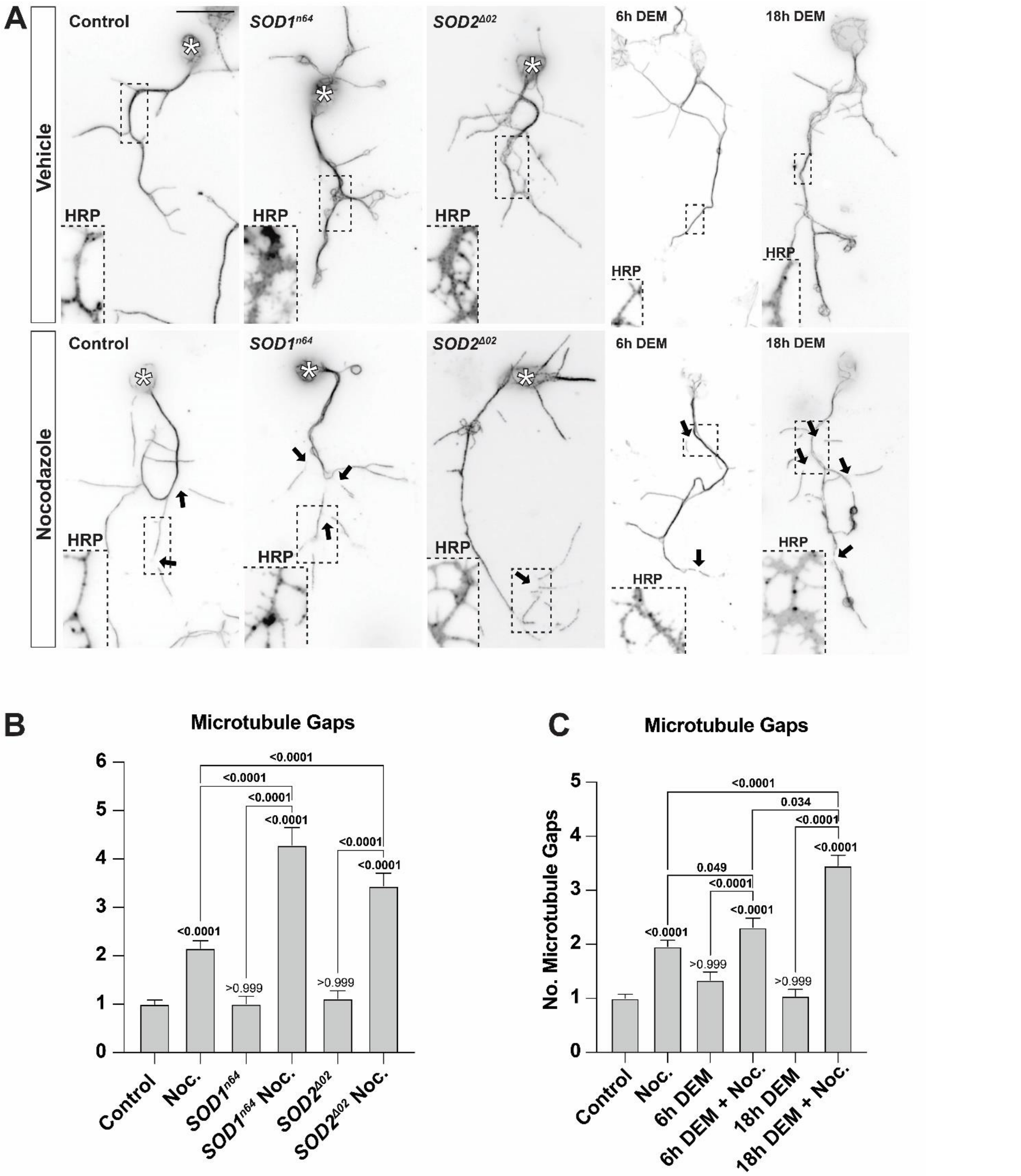


**Figure S2. Conditions of elevated ROS destabilises microtubules in primary neuronal cell cultures.**

(A) Representative images of primary *Drosophila* neurons stained for tubulin and HRP to label microtubules and the cell membrane (only shown in the insets). Neurons of different genotypes: $w^{1118}$ (control) or carrying the $SOD1^{n64}$- or $SOD2^{\Delta 02}$-mutant allele in homozygous, or treated with 100 µM DEM or ethanol at 6 HIV. Cells were treated with 100 µM nocodazole or DMSO at 18 HIV. Black arrows indicate sites of microtubule gaps, induced by nocodazole treatment, indicative of sensitive or unstable microtubules. Dashed boxes show regions of the axon that correspond to each image inset, which show continuous cell membrane using HRP staining. Asterisks label the cell bodies. (B) Quantification of the number of gaps in microtubules per cell; bars represent normalised mean ± SEM. P values are shown above each bar, as assessed by Kruskal-Wallis one-way tests. Data were collated across five independent repeats. A minimum of 60 neuros per condition were evaluated. Scale bars = 10 µm.
