## Supplementary figure 3 for "Oxidative stress promotes axonal atrophy through alterations in microtubules and EB1 function"

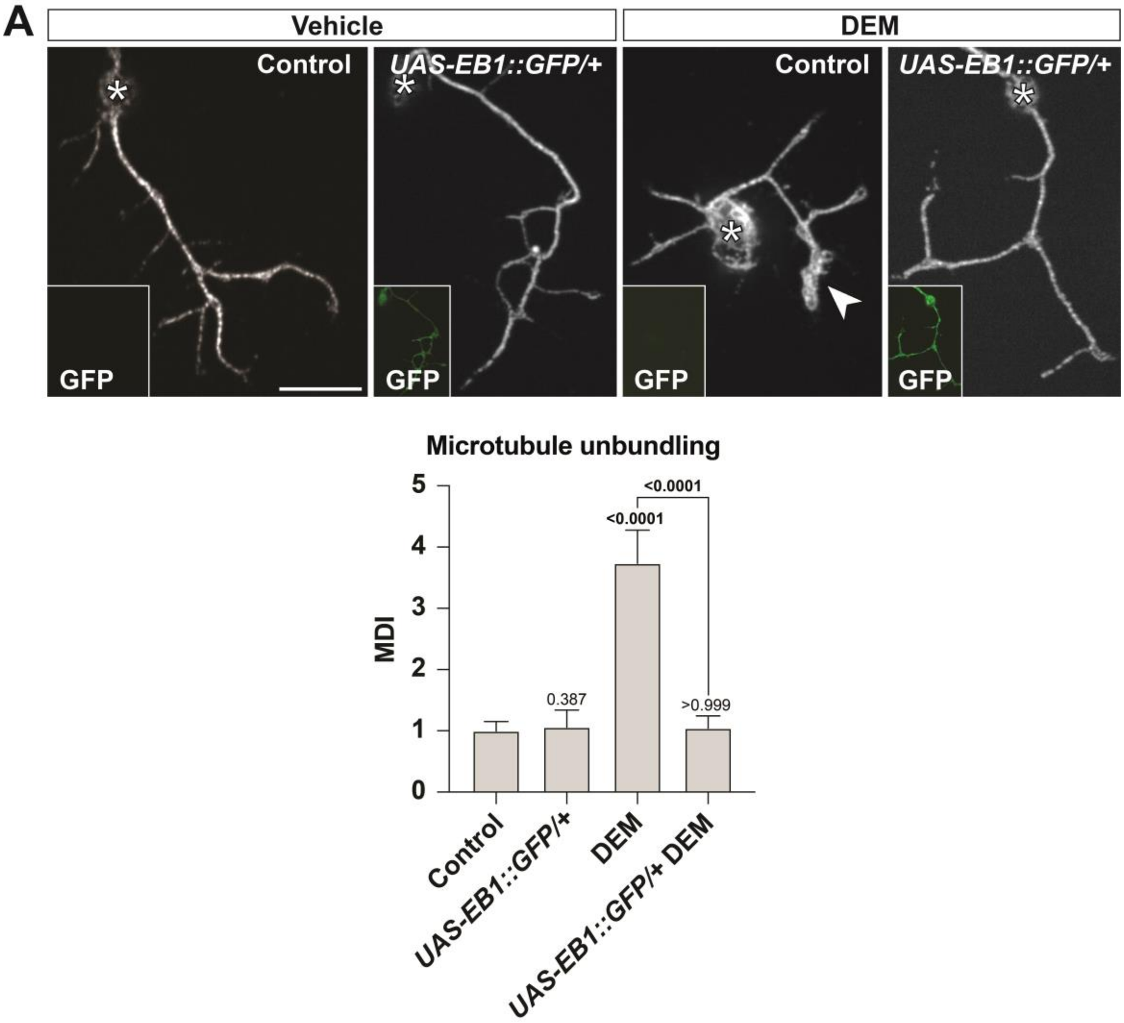


**Figure S3. Overexpression of EB1 prevents DEM-induced microtubule disorganisation**.
(A) Representative images of primary *Drosophila* neurons stained for tubulin. Neurons of different genotypes: *w1118* (control) or overexpressing *EB1::GFP*. Image inset shows whole-cell GFP signal (488 nm excitation). Cells were treated with ethanol or 100 µM DEM at 3 DIV. Asterisks indicate cells bodies and arrow heads indicate regions of microtubule disorganisation. (B) Quantitative analysis of MDI per cell; bars represent normalised mean ± SEM. P values are shown above each bar, as assessed by Kruskal-Wallis one-way tests. Data were collated across 2 independent repeats with 3 individual culture per repeat. A minimum of 80 neuros per condition were quantified. Scale bar = 10 µm.
